# High Resolution Tomographic Analysis of *in vitro* 3D Glioblastoma Tumor Model under Long-Term Drug Treatment

**DOI:** 10.1101/684019

**Authors:** Mehmet S. Ozturk, Vivian K. Lee, Hongyan Zou, Roland H. Friedel, Guohao Dai, Xavier Intes

## Abstract

Glioblastoma multiforme (GBM) is an extremely lethal type of brain tumor as it frequently develops therapeutic resistance over months of chemotherapy cycles. Hence, there is a critical need to provide relevant biological systems to guide the development of new potent personalized drugs but also efficient methodologies that enable personalized prediction of various therapeutic regimens for enhanced patient prognosis. Towards this goal, we report on the development of *i*) an appropriate *in vitro* model that mimics the 3D tumor microenvironment and *ii*) a companion imaging modality that enables to assess this *in vitro* model in its entirety. More precisely, we developed an integrated platform of bio-printing *in vitro* 3D GBM models and mesoscopic imaging to monitor tumor growth and invasion along with long-term drug treatment. The newly-developed *in vitro* 3D model contains tumor spheroids made of patient-derived glioma stem cells with a fluorescent reporter and vascular channels for drug perfusion. The imaging of these thick tissue constructs was performed using our second-Generation Mesoscopic Fluorescence Molecular Tomography (2GMFMT) imaging system which delivered 3D reconstruction of the fluidic channels and the GBM spheroids over the course of pre- and post-drug treatment (up to 70 days). The 2D measurements collected via 2GMFMT was comparable to existing imaging modalities, but 2GMFMT enabled non-sacrificial volumetric monitoring that provided a unique insgiht into the GBM spheroid growth and drug response. Overall, our integrated platform provides customizable *in vitro* model systems combined with an efficient long-term non-sacrificial imaging for the volumetric change of tumor mass, thus has a great potential in profoundly affecting the drug pipeline for a vast array of pathologies as well as for guiding personalized therapeutic regimen.

## Introduction

Glioblastoma multiforme (GBM), a highly invasive malignant brain tumor, carries a dismal prognosis, with a median survival of 14 months [1] and less than 10% of 5-years-survival rate after diagnosis, despite aggressive therapy, including surgery. radiotherapy and chemotherapy [2]. Due to the tumor stemness and heterogeneity of GBM, the tumor cells experience continuous phenotype changes during months of treatment cycles and often develop therapeutic resistance, resulting in a high recurrent rate (>90%), and consequentially high lethality [3]–[5]. Therefore, beyond the necessity to guide the development of new personalized drugs, efficient model systems that enables fast and predictive evaluations of therapeutic efficacy are in critical need. To recreate the tumor responses in experimental settings, either *in vivo* or *in vitro*, a suitable tumor growth environment and long-term culture capabilities are required.

Today, the gold standard to study GBM remains animal models, which typically involve injecting GBM cells into the mouse brain and conducting the histological analysis for different stages of tumor. However, those studies are extremely long, expensive, and suffer from large variability and a poor predictive potential for clinical outcome. *In vitro* approaches including monolayer cell culture [6]–[8] and spheroid/aggregates in suspension [8]–[13] allow high-throughput testing of various therapeutic options. However, these models lack the long-term culture capability and the 3D tumor microenvironment; therefore, have limited abilities to replicate the dynamically-changing tumor behaviors *in vivo*. Maintaining long-term cell viability is also challenging in current *in vitro* models because of the limited territory for proliferation and the difficulties in supplying sufficient nutrient/oxygen transport through diffusion. There is a pressing need for *in vitro* GBM tumor models that can provide sufficient 3D spaces and microenvironments to mimic the invasive tumor behaviors while supporting long-term tissue viability.

Additionally, the assessment of tissue constructs through end-point imaging brings new challenges. 3D model systems are required to be in millimeters-to centimeters-thick to contain a sufficient volume of matrices for long term sustainability but also to mimic the *in vivo* micro-environment. Though, at such scale, tumor tissue structures become more and more turbid due to the increased cell number and the deposition of cell-secreting materials. Hence, traditional imaging technologies such as fluorescence microscopy techniques are limited in assessing these large tissue samples because of limited field of view, shallow depth penetration and potential for photo damage. Although optical imaging techniques have been enabling tumor microenvironment studies for decades, a tradeoff between speed, depth, and resolution is still the inherent limitation [2–4]. Indeed, the optical imaging techniques that are proficient in the mesoscopic regime (a few millimeters depth) over the large field of views are limited to either structural imaging [17] or lack of molecular fluorescence sensitivity (*i*.*e*. photoacoustic imaging). On the other hand, techniques with high molecular sensitivity, *i*.*e*. fluorescence microscopy techniques, suffer from the limited field of view which proportionally increases the imaging time, and the limited imaging depth in turbid tissue structures. Long-term and non-sacrificial imaging that allows saving time and cost is another requirement for efficient and predictive therapy evaluations, as it will ultimately enable high-throughput screening for personalized treatment. Thus, a non-invasive, fast deep tissue imaging technique is required for imaging thick *in vitro* 3D tumor model designed with the proper microenvironment and for longitudinal studies.

In this study, we took a synergistic approach to address the challenges mentioned above by combining a 3D bio-printed tumor-vascular model with our 2nd generation Mesoscopic Fluorescence Molecular Tomography (MFMT) imaging system succeding the first generation [18]. Herein, we demonstrate that our platform enabled up to 70 days of viable tissue culture and 3D volumetric monitoring of patient-derived GBM tumor spheroids during pre- and post-drug treatment periods (Fig.1, top panel). The *in vitro* longitudinal studies span several months, have strict limitation on image acquisition time. The shorter time the viable tissues have been outside of the incubator, the less damages and fluctuations on tissue metabolisms. The required imaging acquisition time has the outmost importance for selecting the suitable modality for the reason. 2GMFMT offers the least stress on cell culture, allows frequent imaging (higher temporal resolution for longitudinal studies) while enabling to image the whole tissue construct (Fig.1, bottom panel).

## Results

### 3D Model Development: *Tumor Cell Responses to Drug Treatment in 2D and 3D Cultures*

The traditional *in vitro* models, 2D cell monolayer (Fig. 2A-B) and 3D spheroid in suspension (Fig. 2C-D), failed to recapitulate tumor invasion features due to the lack of surrounding matrices. In addition, the long-term TMZ treatment was hindered by the overgrowth of cells. 2D cultured GBM cells formed a monolayer of an extremely dense cell population and the layer often peeled off during media change (Supp. Fig. 1), impeding the sample reproducibility. Spheroids cultured in suspension irregularly created satellite cell clumps or 2D growth in the low-attachment well (Fig. 2D), also suppressing long-term reproducibility. The metabolic activities of GBM cells in both conditions were decreased over time with higher TMZ dose. The suspended 3D spheroid (Fig. 2C) presented more sensitive responses to TMZ treatment than 2D monolayer (Fig. 2A), also showing the shrinkage of tumor mass (Fig. 2D). However, both models were not able to capture the re-growth of GBM cells shown in 3D bio-printed model (Fig. 2E).

**Figure 1.**
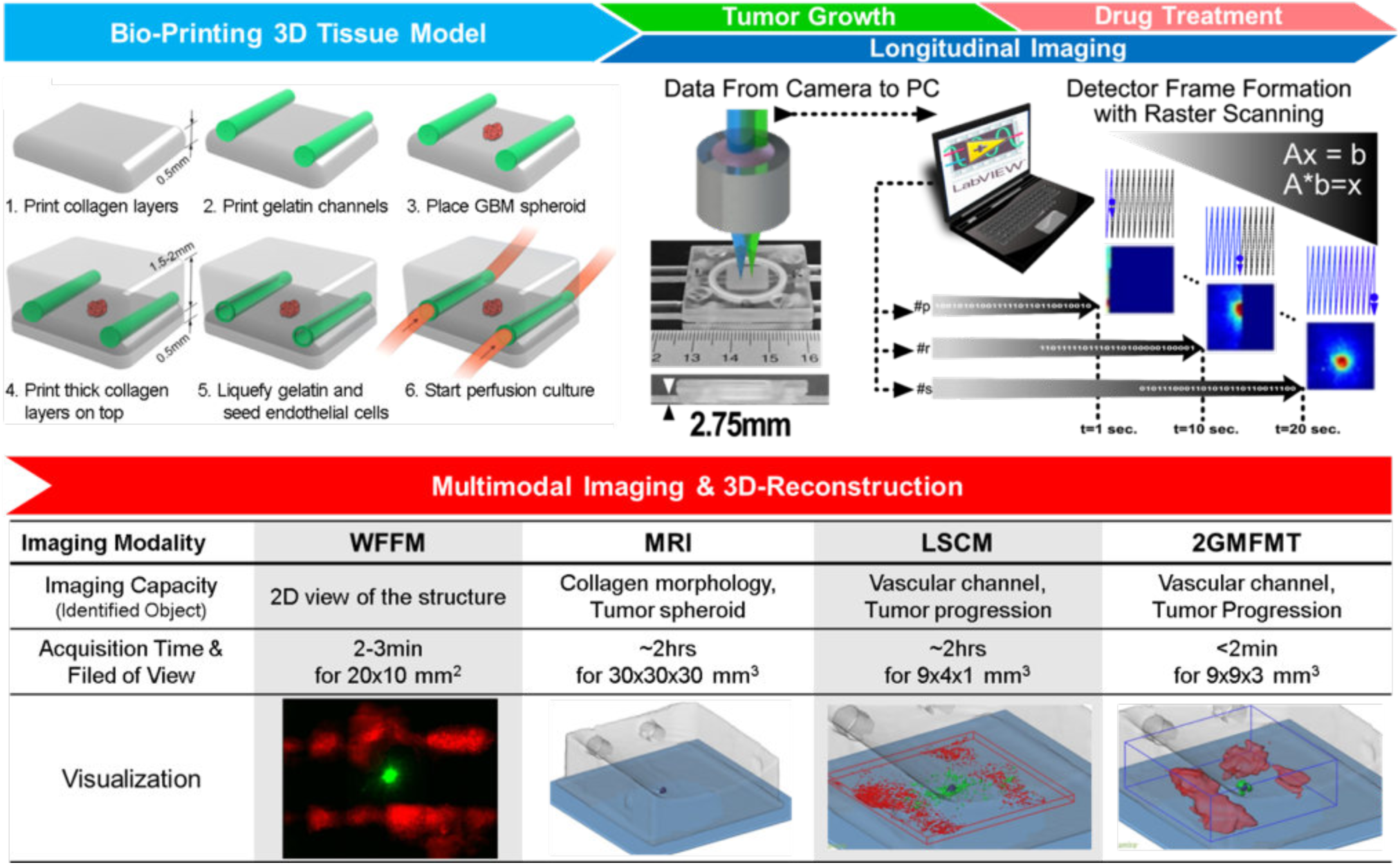
(**Top panel)** The integration of the platform: 3D tumor tissue model and 2GMFMT. The platform enables long-term tissue culture and longitudinal assessment of tumor invasion. **Bio-printing:** 3D tissue containing a GBM spheroid and vascular channels were bio-printed within a plexiglass perfusion chamber and cultured with media perfusion (with or without drug). **Longitudinal Imaging** is conducted through a thick plexiglass for both tumor growth period and drug regimen. Non-invasive imaging was conducted through a transparent plexiglass chamber. De-scanned configuration enabled dense sampling of the target. Each scan point (i.e. p, r, s) represents a pixel in the raw data. As a representative, one detector raw data was shown. A full frame was completed when raster scanning is finished (scan point number, s). Typically, a full field of view scanning is completed in ∼20sec. (**Bottom panel)** shows: The **Multimodal Imaging & 3D Reconstruction** for all potential modalities. 2GMFMT imaging was performed on the model every 3-4 days with 2GMFMT. 2GMFMT presents superior data acquisition time for volumetric assessment of glioblastoma brain tumor in comparison to its counterparts, μMRI and LSCM.

**Figure 2.**
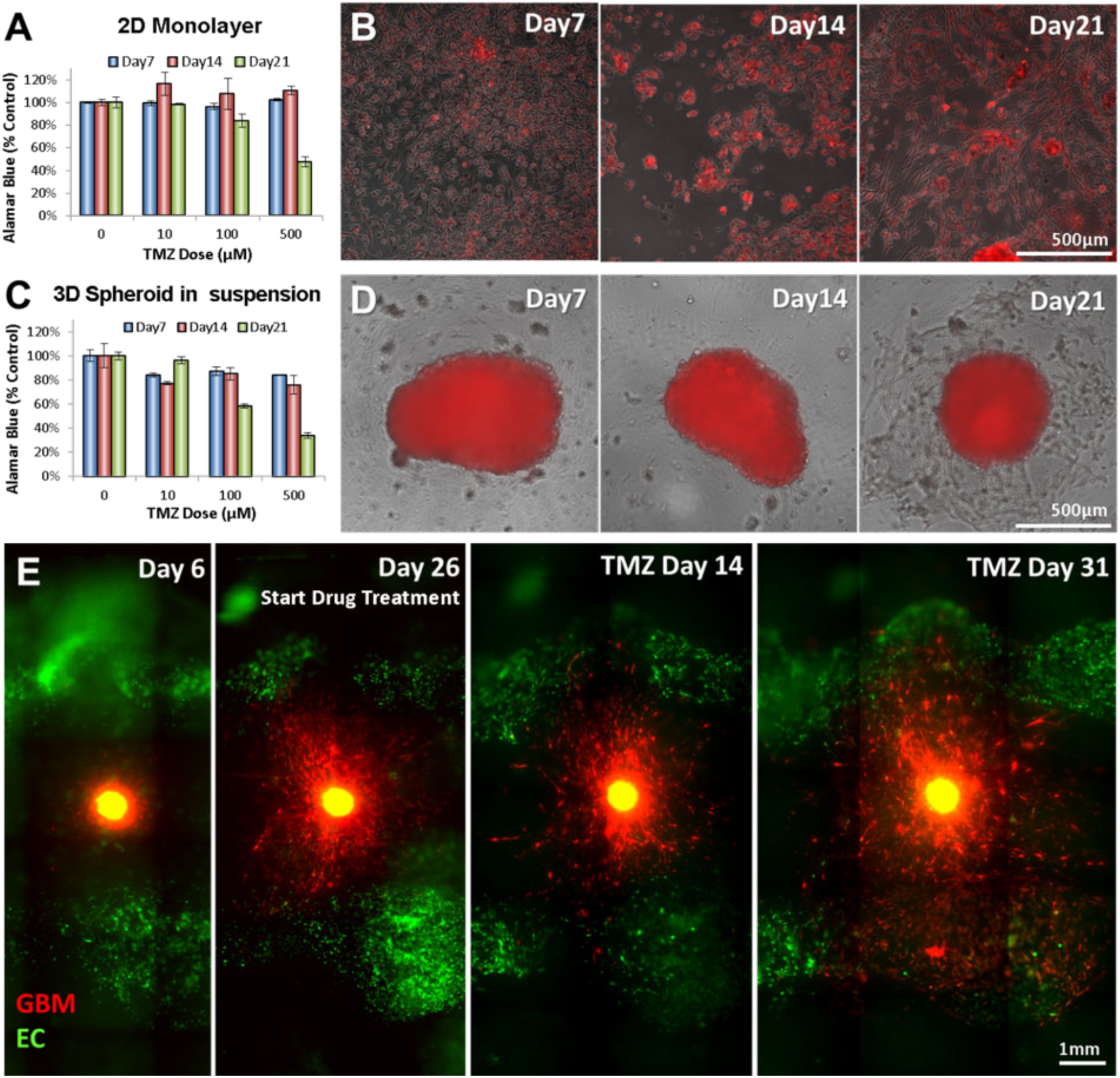
Drug response of GBM cells cultured in different settings. A-D, Alamar blue test results for testing metabolic activities of tumor cells and the overlay of phase contrast and fluorescent images of GBM cells cultured in 2D monolayer (A-B) and 3D suspended spheroid (C-D). (E) Invasive behavior of GBM cells before and after the treatment. SD02 cells from the embedded spheroid aggressively invaded into the surrounding matrix (Day26), regressed and showed shrinkage of tumor core after drug treatment (24 days after drug treatment). However, some GBM cells survived the treatment and resumed its active invasion even with the continuing treatment (31 days after drug treatment). Top-view images (B, D, E) captured using Wide-Field Fluorescence Microscopy (WFFM).

The newly-developed bio-printed 3D tumor-vascular model was cultured under dynamic perfusion for up to 70 days. The structural integrity and the cell viability of the printed tissue were maintained during the culture period, showing the long-term culture and imaging capabilities of our platform. The embedded GBM spheroid began matrix invasion within the first week after the fabrication (Fig. 2E, Day 6). The pre-treatment culture lasted for 3-5 weeks until the GBM cells traveled 1-2 millimeter from the spheroid, invading into the surrounding matrix (Fig. 2E, Day 26). Then, drug-treatment culture started with perfusion of 100µM Temozolomide (TMZ). During the first three weeks of subsequent drug treatment, we observed the regression of individual tumor cells in the invasion area and a slight decrease in the fluorescent intensity (Fig. 2E, 24 Days after Drug Treatment). Despite this regressing tendency, some GBM cells survived the treatment and demonstrated therapeutic resistance [15]. This subset of cells resumed the matrix invasion and proliferation, even with the continuing drug treatment (Fig. 2E, 31 Days after Drug Treatment).

### Resolution Characterization of MFMT

To demonstrate the 3D imaging performances of our 2GMFMT platform (Fig. 3A) in terms of depth penetration and imaging resolution, we designed an agar phantom containing fluorescence polystyrene beads. The beads were embedded inside an agar phantom (z= 1.25mm) which had optical properties similar to brain tissue (μ_s_ ‘=1mm^-1^, μ_a_ =0.02mm^-1^). This phantom was imaged under μMRI (7T Brucker) and Laser Scanning Confocal Microscopy (LSCM – LSM 510 Zeiss) to benchmark the 2GMFMT 3D imaging performances. The optical imaging modalities were conducted through the same side of the chamber that they both experience the same tissue thickness. Results of this comparative imaging study are provided in Fig 3B-D.

**Figure 3.**
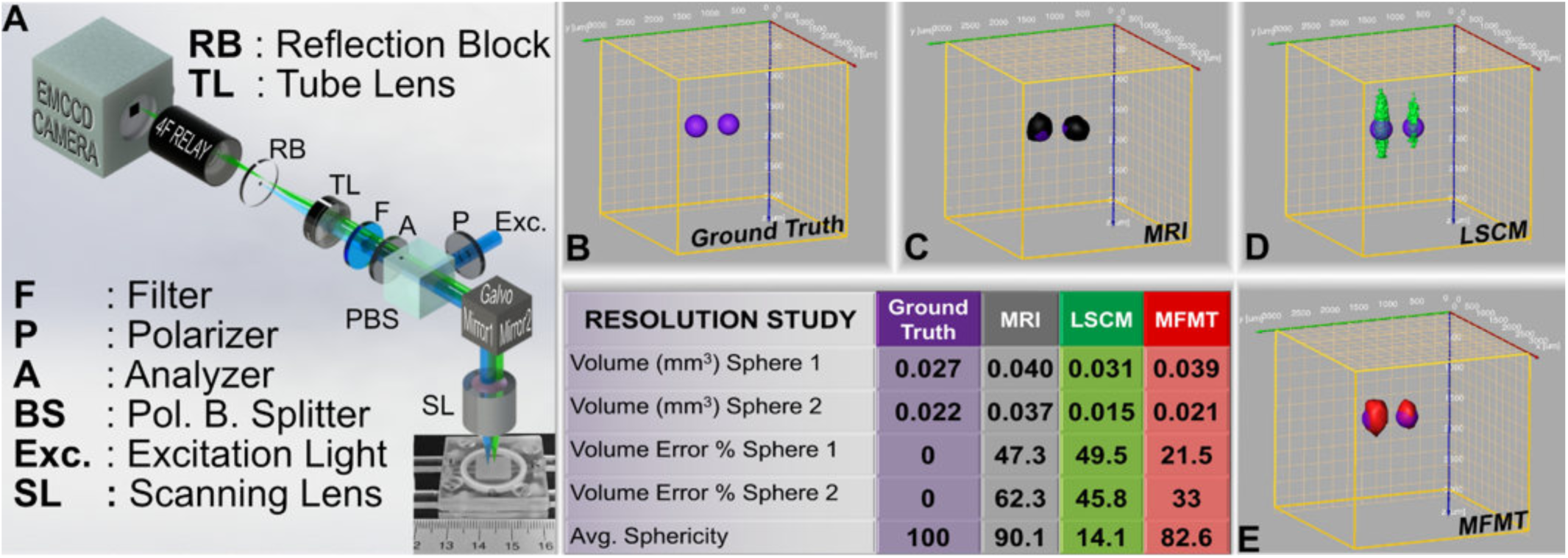
A) 2GMFMT system share a similar pathway for excitation and emission, which enables de-scanned imaging. Polarized (P) excitation signal (Exc) is reflected from Polarizing Beam Splitter (PBS) and raster scanned with 2D Galvo-Mirror and a scanning lens (SL). The emitted light and backscattered excitation light follow the same path back to PBS. The fluorescence emission separated from excitation light with a cross polarizer (A) and a color filter. Tube lens then generate an image plane for Reflection Block (RB), which further reduce the effect of specular reflection (especially in Higher Binning, i.e. Binning>16) and imaged on to the camera. B) A resolution study for fluorescence beads was selected as a ground truth. *μ*MRI (C), LSCM (D) and 2GMGMT (E) competed against each other. Performance criterion table of volume error, and sphericity showed that 2GMFMT delivered superior volume accuracy and competing sphericity.

To assess objectively the performances of 2GMFMT versus the two conventional imaging modalities selected, we computed image-driven quantitative metrics using a commercial software (Amira, FEI Inc.). These metrics include volume estimate as well as sphericity. Our sphericity metric reports on the accuracy of the reconstructed spherical shape and was computed following a similar approach as used in image processing literature [4, 5]:

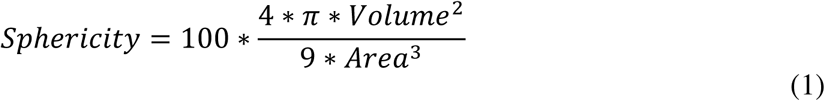

A perfect sphere yields a *Sphericity* score of 100 and lower scores indicates a loss of accuracy in retrieving the spherical bead shape. Overall, 2GMFMT resolved two spheroids either better or competitive performance metrics in comparison to conventional imaging modalities.

Our results indicated that the 2GMFMT delivered the smallest volume error for both spheres. LSCM yield the lowest accuracy in shape despite the highest spatial sampling among the three modalities. Thanks to the strong refractive index difference between the polystyrene spheres and agar phantom, μMRI yield the highest shape accuracy (∼90%), which is followed by 2GMFMT with ∼83%. On volume estimation, 2GMFMT delivered the lowest error with ∼33% (avg.) while μMRI and LSCM had much higher average error, ∼62% and ∼43%, respectively (Fig.3). Though it is important to note that 2GMFMT data acquisition took much less (<2min) than μMRI (∼2hrs.) and LSCM(∼3min).

### Tissue Culture Imaging

We analyzed the image end points in two folds: 2D intensity–area comparison and 3D reconstruction-volume comparison.

#### Area-Based Comparison

To further validate the potential of our combined bio-printing and imaging platform, we compared raw fluorescence measurements obtained via Wide-Field Fluorescence Microscopy (WFFM) and our 2GMFMT system (Fig. 4A-B). In clinical settings, the treatment efficacy and the disease recurrence are typically assessed via medical image-based endpoints including the tumor diameter(s) (*i*.*e*. cross sectional area) and tumor volume for the drug response assessment [21]. Since WFFM does not have 3D capabilities and its measurements were limited to 2D, we performed area-based comparisons using the epi-fluorescence microscopy equivalent of 2GMFMT measurements (i.e. zero source-detector separation data). We picked tumor H for demonstration purpose and rest of the comparison can be found in the supplementary figures (4-6), presenting the corresponding trends between WFFM and 2GMFMT. In both modalities, we observed a decreased fluorescent intensity in the tumor core area and the spreading of peripheral low-intensity area which represents individual tumor cell invasions (invasion area). Even though 2GMFMT could not provide a single cell-scale resolution as WFFM, it successfully captured the changes in intensity and tumor area with 200µm lateral resolution. To further refine the utility of such 2D measurements, the fluorescence signals associated to the spheroid (core) and invasion components of the tumor model were retrieved via image segmentation (Fig. 4E-F). The core was represented by the full width half maximum (FWHM) intensity, and the invasion area was depicted by the intensity profile from FWHM to 10% of the maximum. The intensity values below 10% were regarded as background signal and discarded. The longitudinal quantitative estimation of these two 2D measurements is reported in Fig. 4C-D. The timeline divided into a growth period (green area: prior to drug treatment; pre-treatment) and a drug treatment period (red area: post-treatment). Overall, the trend in the rate of changes in core shows a difference as the tumor was continued to receive TMZ drug. But the invasion areas were similar throughout the >2 months imaging.

**Figure 4.**
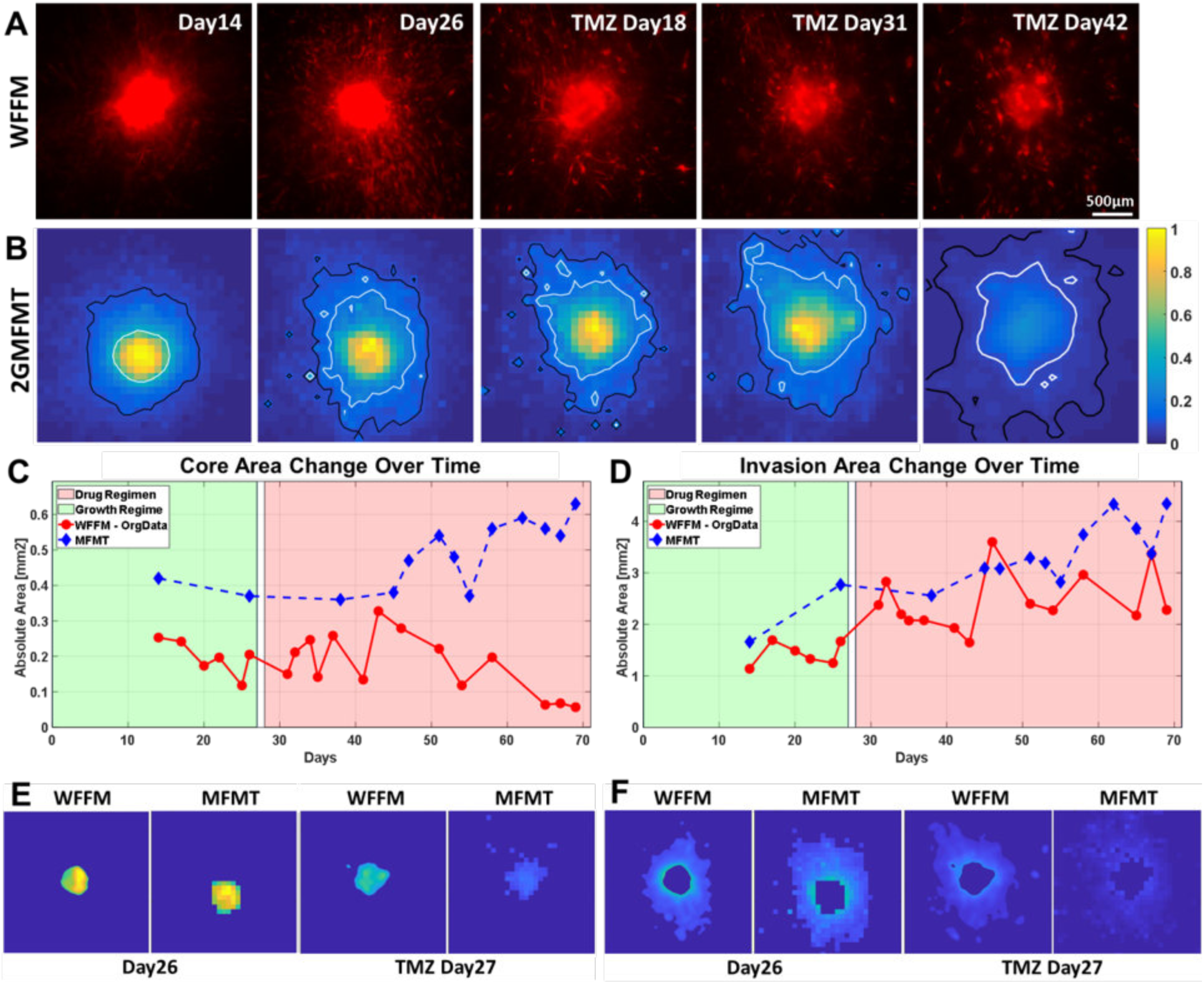
Longitudinal intensity assessment of Glioblastoma Brain Tumor H with A) Wide Field Fluorescence Microscopy (WFFM) which depicts the cellular invasion B) Epifluorescence equivalent raw data from 2GMFMT shows the diffused intensity signal from tumor cells. The 50% maximum intensity isoline (white) delineate the tumor core area and 10% maximum intensity isoline (black) shows the invasion area. Core Area (C) and Invasion Area (D) values are shown for all data points and also representative data points are shown pictorially. Especially Invasion area followed the same trend across modalities. (E) shows the segmented core areas from WFFM and MFMT 2D images for initial stage (Day 26) and regressed stage (TMZ regimen at Day 27). (F) shows the segmented invasion areas WFFM and MFMT 2D images for initial stage (Day 26) and regressed stage (TMZ regimen at Day 27).

#### Volume-Based Comparison

2D measurements such as cross-sectional estimates provide only a partial view of the tumor model. Hence, clinically, tumor treatment efficacy is assessed by volume estimation through μMRI where the volume change is expected to represent the overall tumor response against therapies. *In vitro*, the tumor volume could be non-invasively estimated via 3D imaging of the fluorescence signals associated with the tumor-tagged reporter gene. Advanced fluorescence imaging modalities such as Confocal or Multiphoton can be employed for the purpose when the sample is relatively transparent and thin (a few hundred micrometers in thickness). However, our 3D tumor tissue model system that consists of a 3-5 mm thick turbid construct and a sealed biochamber is 8-10 mm in total thickness, thus not amenable to Confocal or Multiphoton.

In order to demonstrate the utility of our mesoscopic imaging system on such models, the tumor models were imaged sequentially on μMRI, LSCM and 2GMFMT up to 70 days including tumor growth regime and drug regimen. The 3D visualization of the tumor model (red) and the printed perfused channel (green) are provided in Fig. 5A. 3D reconstruction of 2GMFMT provides diverse aspects of information/knowledge on the volumetric change of GBM tumor. The tumor spheroid maintained a spherical shape during the growth period (Day 26) and the drug treatment (Day 63: Drug Day 37), then transformed to the flatter shape and invade into surrounding area (Fig. 5A). These observations from the volume-based analysis of 2GMFMT correspond with our previous findings on drug-responsive tumor behavior (Fig. 2E). While 2GMFMT successfully captured the volumetric information of tumor mass, LSCM partially detected the bottom hemisphere of tumor spheroids (Fig. 5A, LSCM, Day26). As the tissue structure became more opaque throughout the culture period, it became more difficult to retrieve fluorescent signals using LSCM (Fig. 5A, LSCM, Day 63: Drug Day 37). The LSCM imaging acquisition was conducted through the bottom part of the collagen matrix (spheroids are within ∼500µm deep) whereas the 2GMFMT imaging was conducted through the top part of the collagen (spheroid ∼1-2mm deep), which is far more challenging configuration.

**Figure 5.**
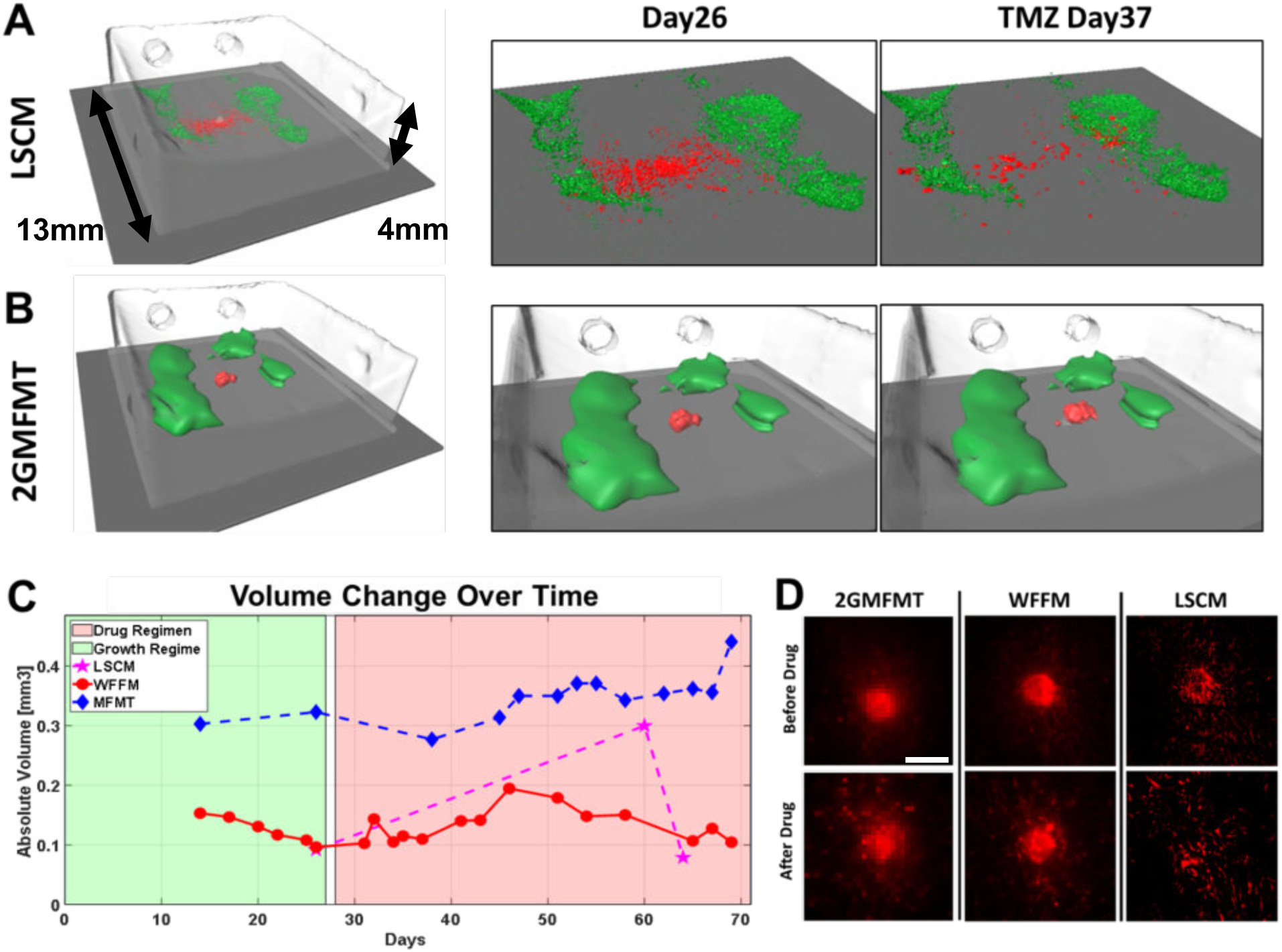
Longitudinal Volumetric Assessment of Glioblastoma Brain Tumor was demonstrated by comparing Laser Scanning Confocal Microscopy (LSCM) and 2GMFMT (A-B). μMRI provided collagen structure and was used as a co-registration land mark. Vascular construct (Green) and GBM (Red) is visualized by both LSCM and 2GMFMT. (C) LSCM and 2GMFMT showed a similarity in volume variation trend with different rate of change. (D) Maximum intensity projections from LSCM and 2GMFMT were compared with LSCM images (Scale bar, 500 µm).

3D tumor volume was estimated using the measurements from the three different imaging modalities (Fig 5B). For 2GMFMT, the tumor volume was calculated by the voxel numbers. The results show a gradual volume change trend with a slight increase during growth period (Fig. 5B, green area), a brief stagnation of growth when the treatment began, and a continuous expansion through the rest of treatment cycle (Fig. 5B, red area). For WFFM, the volume was estimated using 2D cross-sectional area measurements with an assumption that tumor spheroids are relatively sphered in shape. Due to the nature of the estimation process, the final volume calculation can be hugely affected by minor deviations such as an irregular/asymmetrical form of tumor mass or a couple of tumor cells that travel farther than the majority. This caused the discrepancy in volume calculation at the later stage of drug treatment (Fig. 5C). When GBM spheroids maintained a spherical shape, the volume calculations from WFFM and MFMT were well-corresponding (Supp. Fig. 4-6D). In the case of LSCM, it was impossible to obtain meaningful volume data as no signals were captured when imaging with LSCM at depth over half a millimeter. 3D estimation of the tumor volume via μMRI measurement was possible only at the initial stages when the contrast between the spheroid and the collagen matrix is large enough. μMRI failed to detect the later tumor expansion led by GBM cell invasion and migration.

## Discussion

Clinically relevant biological systems with predictive potential allow to investigate patient-specific tumor behavior and to tailor personalized therapeutic strategy, thus will improve GBM patient prognosis. To enable such goals *in vitro*, experimental models need to satisfy multiple requirements: *i*) the use of patient-derived GBM cells that can replicate the dynamic tumor behaviors exhibited *in vivo*; *ii*) controllable 3D microenvironment that provides space and stimuli for 3D tumor invasion which is a marked feature of GBMs; *iii*) long-term culture and imaging capability; and *iv*) non-sacrificial non-invasive imaging modality that can longitudinally detect tumor cells within thick and opaque tissues. Herein, we report on the development and characterization of a study platform that meets these requirements. The integrated platform of bio-printed tumor-vascular model and 2GMFMT supported a faster acquisition time compared to the traditional modalities, minimized the tissue damage, thus enabled successful long-term monitoring of tumor invasion with higher temporal resolution. The volume investigation was conducted pre- and post-drug treatment, showing the potential of our system for the creation of patient-specific tumor model and its application for testing various treatment regimens (*e*.*g*. chemotherapy, radiotherapy, combinations of multiple treatments).

On the bio-printing side, the 3D constructed model allows replicating 3D environment for the physiological tumor invasion and the anti-treatment responses. The incorporation of tumor spheroid within 3D matrix not only allows the GBM invasion but also increases the lifespan of tumor spheroid. In the tumor spheroids cultured in suspension (Fig. 2C-D, Supp. Fig. 2), substantial cell death begins after 2-3 weeks of culture, due to the diffusion limit of nutrient and oxygen through the dense cell aggregate. Unlike the suspension culture, the spheroids within 3D matrix present organoid-like behaviors [14, 15], presenting an extended lifespan and stratified cell layers. The fluorescent intensity of core area was reduced after the drug treatment, indicating significant cell death in the area and possibly a formation of necrotic tumor core (Fig. 4E). The tumor cells in the core region (inner layer) and the surround region (outer layer) showed distinctive morphologies, proliferations, and invasions (Fig. 4E-F).

The major advantage of our system over other in vitro models is that the vasculature is constantly perfused and resides in a biologically relevant matrix to allow native cell-ECM remodeling and efficient drug delivery. The customization capability of tumor environments is also a strength of our system. Bio-printing technology allows incorporating multiple cell types, ECMs, and soluble chemical in desired 3D patterns. These features broaden the application of our system. For example, a 3D model that more closely mimics brain environment can be established by incorporating astrocytes and glial cells around the tumor spheroid. This system presents a unique experimental tool to address fundamental questions of tumor vascular biology, which would be difficult or impossible to resolve using conventional *in vivo* or *in vitro* models.

Imaging such models in their entirety and longitudinally is very challenging through existing commercial imaging modalities. Hence, we integrated the bio-printing approach with an advanced mesoscopic optical imaging modality, 2GMFMT. To demonstrate the utility and performances of this novel molecular imaging methodology, we performed: *i*) Data fidelity check (WFFM and 2GMFMT) and *ii*) Volumetric assessment across modalities (μMRI, LSCM, and 2GMFMT). In summary, 2GMFMT can collect the same measurements that can be gained by current non-sacrificial imaging modalities, such as tumor area and fluorescent intensity change, with an equivalent or higher sensitivity. In addition, it has a superior capability in the 3D visualization and the volumetric monitoring of GBM tumor growth within thick and opaque tissue constructs. The intensity profile and cross-sectional area, measured by 2GMFMT, was comparable to WFFM (Fig. 4). For the volumetric assessment, μMRI gave accurate data on collagen density but could not discern spheroids from the collagen scaffold in the later culture stages, which was in a necessity for validation purpose. LSCM, on the other hand, failed to collect accurate volume information since it is incapable of capturing the signal from the entire volume, deep in the collagen matrix. Overall, the 2GMFMT provides accurate sphericity, fluorescent intensity, and volume. Other microscopic techniques such as light-sheet microscope [24] are valuable here, but it requires sample fixation and/or clearance for optical transparency. 2GMFMT, on the other hand, can be applied to live samples without fixation or any chemical pre-treatment. For those reasons, we believe that the integrated platform of 3D bio-printing and 2GMFMT has a potential to be a valuable addition to longitudinal tissue imaging toolbox.

It is also important to note that the imaging of biological samples in their biochamber leads to additional challenges. Widely-used imaging technologies for tumor studies, μMRI and LSCM, have a long time-requirements for *in vitro* and *in vivo* studies. Herein, μMRI and LSCM imaging session required ∼2hours to complete. This puts an immense pressure on the viability of cells. Conversely, 2GMFMT system allows full volumetric data acquisition in as quick as 20 seconds. Thanks to high density sampling through 2D detector array of CCD camera, 2GMFMT can deliver high resolution reconstruction (100-200μm) over a large field of view (up to 1cm2) up to 5mm depth. Those key features provide a big relief for longitudinal imaging of tissue samples, which need to be taken out of the incubator and put back as soon as possible after the imaging acquisition. By definition, those tissue constructs host highly scattering tissue samples and should be kept in a controlled environment (*i*.*e*. incubator). Access to an imaging modality that can complete data acquisition for volumetric imaging over a large field of view in less than a minute will immensely increase the survival rate of the cells in tissue construct.

In this study, our integrated platform provides customizable *in vitro* model systems combined with an efficient long-term non-sacrificial imaging for the volumetric change of tumor mass, thus has a great potential in evaluating personalized therapeutic options. The next generation *in vitro* models, in synergy with sensitive multiplexed molecular imaging techniques, may offer the unique ability to better understand and characterize GBM biology, guide development of novel therapies, and provide analytical predictive metrics of therapy efficacy for tailoring personalized treatment.

## Materials and Methods

### Cell culture and hydrogel preparation

Human umbilical vein endothelial cells (HUVECs; mCherry-transfected) were cultured at 37°C in 5% CO2 in EGM®-2 Endothelial Cell Growth Medium-2 (Lonza). Patient-derived glioblastoma multiform (GBM; EGFP-transfected) cells were cultured on laminin-coated tissue culture flask in NeuroCult™ NS-A proliferation media for human (STEMCELL Technologies). Culture media was changed every two days. For the cell seeding on bio-printed channels, HUVECs were harvested using 0.25% Trypsin-EDTA, and then maintained as cell suspensions on ice until ready to be seeded. To create GBM spheroid, 1,000 – 5,000 GBM cells were plated into Corning® Spheroid Microplate, then cultured for 7-14 days until the spheroids reach the desired diameter (> 400µm). Collagen hydrogel precursor (3.0 mg/mL; Rat tail, type I; Corning) was used as a main scaffold material for bio-printing. Gelatin from porcine skin (10%; Sigma-Aldrich) was used as a sacrificial material to create fluidic channels.

### Bio-printing of vascular-GBM model

Two fluidic vascular channels were created on top of printed collagen layers using 10% gelatin as a sacrificial material (Fig. 1 Bio-printing, step 1-2) [8–11]. GBM spheroid was placed on the printed collagen layer, in between two gelatin channels (Fig. 1 Bio-printing, step 3). Excessive media around the spheroid was removed and a small amount of collagen I was added to fix the spheroid location. More collagen layers were printed on top of the gelatin channels and GBM spheroid (Fig. 1 Bio-printing, step 4). The whole structure was then incubated for 20-30 minutes to liquefy gelatin and obtain fluidic channels. HUVECs in suspension were injected into the channels (seeding density: 8 million cells/mL) to create cell lining on the inner channel surface (Fig. 1 Bio-printing, step 5). The entire construct was printed in a flow chamber which allows stable, long-term perfusion (Fig. 1). The construct was cultured with EGM-2 media for up to 70 days at 37°C in 5% CO2. The culture media was changed 3-4 times a day through the vascular channel.

### Drug treatment

GBM cells were plated in 96-well tissue culture plate (2D monolayer culture condition) or Corning® Spheroid Microplate (suspended 3D spheroid culture condition). The cells were cultured in NeuroCult™ NS-A proliferation media until they formed a confluent monolayer or spheroid with ∼500µm of diameter. Then, the cultured media was switched to EGM-2 media with varying concentrations of Temozolomide (concentrations = 0, 10, 100, 1000µM; Sigma Aldrich). In bio-printed 3D tissue model, drug treatment had begun by adding Temozolomide (final concentration = 100µM; Sigma Aldrich) to culture media when the GBM invasion distance reached 1-2 mm.

### Metabolic activity assay

Alamar Blue® reagent was used to quantify the metabolic activity level of GBM cells after drug treatment. After 7, 14, 21 days of drug treatment, EGM-2 media containing 10% Alamar Blue was added to 96-well tissue culture plate (2D condition) and spheroid microplate (suspended 3D spheroid condition). The cells were incubated with Alamar Blue for 4 hours, then the media was collected. The fluorescent intensity of media was measured using Tecan Infinite 200 PRO multimode reader (excitation = 545nm, emission = 590nm).

### Multimodal Imaging Process

Four GBM tumors from two different cell lines were monitored up to 72 days. The longitudinal process includes two segments: a growth period and a drug regimen for the GBMs. The state of the GBMs was assessed by non-concurrent imaging with Widefield Fluorescence Microscopy (WFFM), LSCM, μMRI and 2GMFMT. Our goal is to monitor the cellular invasion and volumetric change of the GBM spheroids and to show that 2GMFMT delivers the most efficient and comprehensive information about the overall structure.

The imaging orientation is shown for LSCM and 2GMFMT in Fig.1. The main reason we chose this orientation was to conduct our validation study. We placed the GBM and vascular channels close to bottom plexiglass (< 500µm) so that LSCM could work in its best condition while 2GMFMT collected the data from the top through ∼3mm thick plexiglass and a few millimeters of collagen.

The longitudinal images were acquired pre- and post-administration of a clinically approved drug (Temozolomide; TMZ; 50-100µM) to replicate the progression of the disease and the clinical protocol where imaging was carried out before initiating the treatment and after treatment ends. Typical acquisition times varied across the imaging modalities: ∼2 minutes for WFFM, 1-2 hours for LSCM, and ∼2 hours for μMRI.

### 2GMFMT Imaging

The optical diagram of our 2nd generation MFMT system is shown in Fig. 3. Briefly, the optical path starts with introducing an excitation light (Ex) into the system through a linear polarizer (P). A Polarizing Beam Splitter (PBS) reflects ∼90% of the S-polarized light onto a Galvanometer Mirror pair (GM). The GM controls the scanning area and dwell time for each excitation point through a Scan Lens (SL). The backscattered light is collected by the same SL and filtered by the PBS. The PBS let pass ∼90% transmission of P-polarized emission and minimize the specular reflection of S-polarized excitation. After the PBS, the backscattered light is further filtered by another polarizer (A – to additionally reduce specular reflection) and then spectrally filtered using the appropriate interference filter (F). As this de-scanned configuration collects light exiting the tissue, 0.6-1.8mm away from the illumination spot, the signals acquired exhibit a large dynamical range. To mitigate this drawback, we introduce a custom-made reflection block (RB) right before relaying the signal onto the camera, effectively blocking the light originating at the same location as the illumination spot. RB ensures an adequate dynamical range and SNR for distal detectors. The light is then collected by an EMCCD (iXonEM+ DU-897 back illuminated, Andor Technology).

### Data Acquisition

The relative positions of source and detector locations as imaged on the specimen by our system are shown in Fig. 4, where the blue dot represents the detectors, red dots represent the source positions. Source raster scans over the target area (Gray Area). Each source point corresponds to a pixel on image plane and as the raster scan completed, full detector frames are formed. This depiction renders an acquisition mode with 2×2 binning, leading to a total of 256×256 detectors. Of those detectors, we selected an appropriate source-detector separation (∼0.6 mm) for 49 detectors in a square grid formation. The center detector delivers the wide-field equivalent image. The source and detectors are moving together as indicated by the red arrow and the source is always in the center of the detector array. The software-controlled raster scanning step size and dwell times are typically set to 200 µm and 20 ms, respectively. The typical FOV of imaging is 6.2×6.2 mm2, leads to scanning a total of 961 points [12] in ∼20sec.

## Reconstruction Algorithm

### The Forward Model

The Radiative Transport Equation was solved (Green function, *G*_*i*_) through CPU based Monte Carlo (MC) simulation. The simulation is computational heavy process due to large number of source-detector pairs (48) and dense spatial sampling (31×31=961). Traditionally, full sensitivity matrix requires a total number of 48×961 = 46,128 simulation. However, thanks to the efficient adjoint formulation [29], we simulated 1 matrix for source (i.e. source, *G*_*exc*_) and 48 matrix for each detector (i.e. *G*_*det*_). Then the multiplication of these two green functions delivered the photon propagation model for 48 source-detector pairs. Then the matrices were populated over the sampling plane (over 961 source positions), which represents the detector readings for all source positions. The simulation was run 10^6^ photons for matching optical properties (*µ*_s_ ‘= 1mm^-1^, *µ*_a_ = 0,02mm^-1^). In a personal computer (Intel XXX, 64 GB Memory), a simulation for a kernel matrix (1 source – 48 detector) took 1-1,5 hour, depending on the number of source locations. One can seamlessly speed-up the process by switching to a GPU based MC.

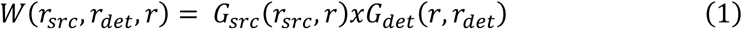

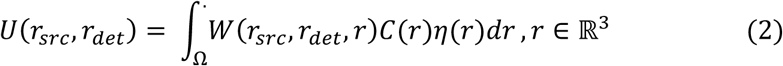

### The Inverse problem

Diffuse nature of the light imposes an ill-posedness to the problem and having an epi-configuration hinders the independent measurements. Thus, sorting out the useful information becomes the key under these conditions [30]. The unknown in this study perceived as composite value of *x*(*r*) = *C*(*r*) * *η*(*r*). By knowing the extinction coefficient and quantum yield of the fluorophore, one can extract the exact concentration [31]. For this paper, it is sufficed to get relative distribution, *x*(*r*). Solving the equation above for the spatial distribution of fluorophore concentration requires an iterative process and up to date number of methods were proposed. Here, we utilize an L_p_-norm regularization scheme, expressed as the following optimization problem [32]

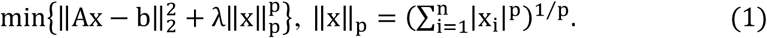

where A is the sensitivity matrix, generated by MC, b is the measurement vector (Supp.3). For this study, we utilized the *L*_*1*_ *-norm* regularization. A spatially variant regularization parameter, *λ*(*r*), determined through the L-curve method [33], helped alleviating the ill-posedness of the problem.

### Multimodal Imaging and Image Registration

Our custom-made plexiglass perfusion chamber enabled a multimodal imaging without disturbing the media. Our imaging procedure began with μMRI imaging and Laser Scanning Confocal Microscopy (LSCM) and concluded with MFMT data acquisition. Due to the long acquisition time in μMRI and LSCM and to avoid the deprivation of cells from oxygen and nutrition, our experiments include a smaller number of data points for those modalities. Data points from those imaging modalities helped us to reveal the trend for volumetric change while MFMT delivered more detailed variation in volume change.

### MRI Imaging

We applied two different protocols (i) Polystyrene & Agar Phantom and (ii) Glioblastoma & Collagen sample due to their different composition and water content. The μMRI protocol agar phantom (Bruker 7T μMRI) used an *Echo time* of 72 msec with *Rare factor* of 12 and *Repetition rate* of 1300 msec for 100 μm isotropic resolution over 2×2×2 *cm*^3^ volume. The protocol for collagen structure used 80 msec *Echo time* with 12 *Rare factor* and 1200 msec *Repetition time*. The imaging volume was 3×3×3*cm*^3^ with isotropic cubic voxels of 150 μm. Data acquisition took 2 hours and 18minutes. μMRI images provided both the location of spheroids and the shape of the 3D printed collagen. To get the collagen morphology within the plexiglass chamber, we used 80msec *Echo time* with 12 *Rare factor*. The top surface of the collagen had a concave curvature and thickness get smaller by time. This information played a key role adjusting the focal plane for MFMT and co-registering the 3D reconstruction over the collagen phantom.

### Laser Scanning Confocal Microscopy Imaging

We used a conventional Confocal Microscope (LSM 510, Zeiss) for both polystyrene bead and glioblastoma imaging. For resolution study, we used 543nm HeNe laser and HFT UV/488/543/633 dichroic mirror BP 565-615 filter pair to match the excitation spectra of RFP polystyrene beads (Cospheric Inc., USA). For vascular channel and GBM spheroid imaging, we used the 488 Argon and 543 HeNe laser, respectively. BP 500-530 IR and BP 565-615 IR were used on the illumination path. For both excitation/emission configurations, we used Plan-Neofluar 10×/0.3 objective, to take advantage of long working distance (5.2mm) due to thick (2.75mm) plexiglass optical window. Fluorophore bead imaging was completed with single data acquisition where both beads fit in the field of view, 900×900μm2 (128×128 pixels) with 7×7×30μm3voxelization for which data acquisition took 1sec./layer for 10 frame averaging. In total, total volume data acquisition took 80 sec. Perfusion chamber imaging required mosaicking due to large interrogation area, minimum 3×9 tiles, which typically took 1-2 hours to complete the data set for 1.75×1.75×20μm3 (512×512 pixels) voxelization of 900×900μm with averaging over 10 frames.

## Acknowledgment

Authors thank Scott McCallum for helping to develop MRI protocol and Sergey Pryshchep for his support setting up parameters for Laser Scan Fluorescence Microscopy.

## Author contributions

M.S.O., V.K.L., X.I. & G.D. conceived the original idea. M.S.O, V.K.L., X.I. & G.D. designed the research study. V.K.L. conducted the cell culture study, built tissue printing platform and acquired the WFFM images. M.S.O. built 2GMFMT system, acquired the LSCM, MRI data and reconstructed 3D images. M.S.O., V.L., X.I., G.D. interpreted the results. M.S.O., V.L. wrote the manuscript. X.I. & G.D. revised and edited the manuscript. All authors discussed the conclusions and commented on the manuscript.

## Supplementary Figures

**Supplementary Figure 1.**
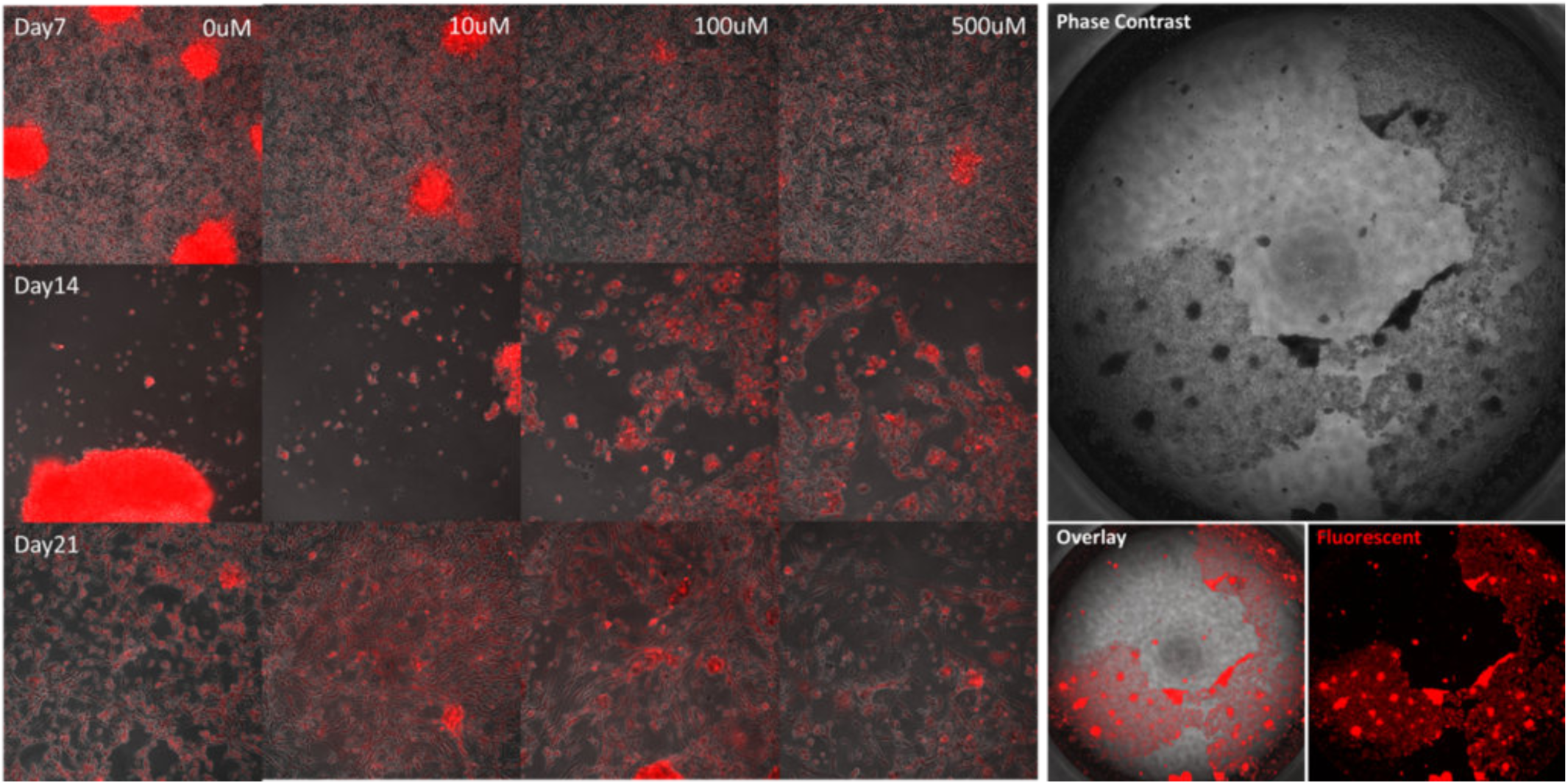
2D monolayer culture of GBM cells. (left panel) GBM cells cultured on tissue culture plate for 21 days with various TMZ dose (0-500uM). Over-populate cell layers contracted, forming clumps and began to peel off after 14 days of culture. (right panel) One example of cell layer peeling-off. Phase contrast and red fluorescent imaging.

**Supplementary Figure 2.**
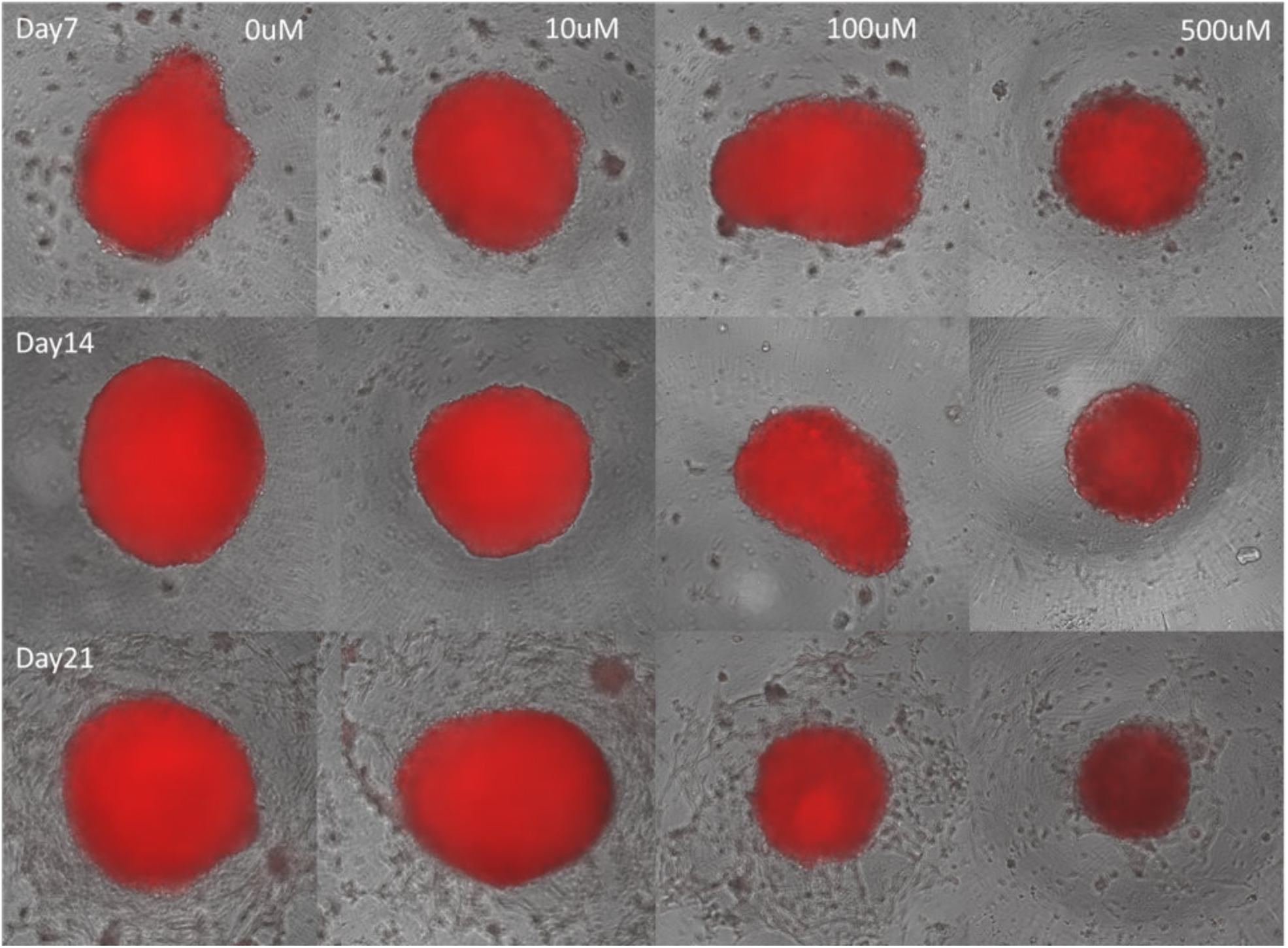
Suspension culture of 3D GBM spheroids. GBM spheroids were cultured in suspension for 21 days with various TMZ dose (0-500uM). With higher TMZ dose (>100uM), we observed the shrinkage of spheroid mass and the decrease in fluorescent intensity. GBM cells began to attach and proliferate on the surface of low-attachment spheroid culture plates after 14 days of culture.

**Supplementary Figure 3.**
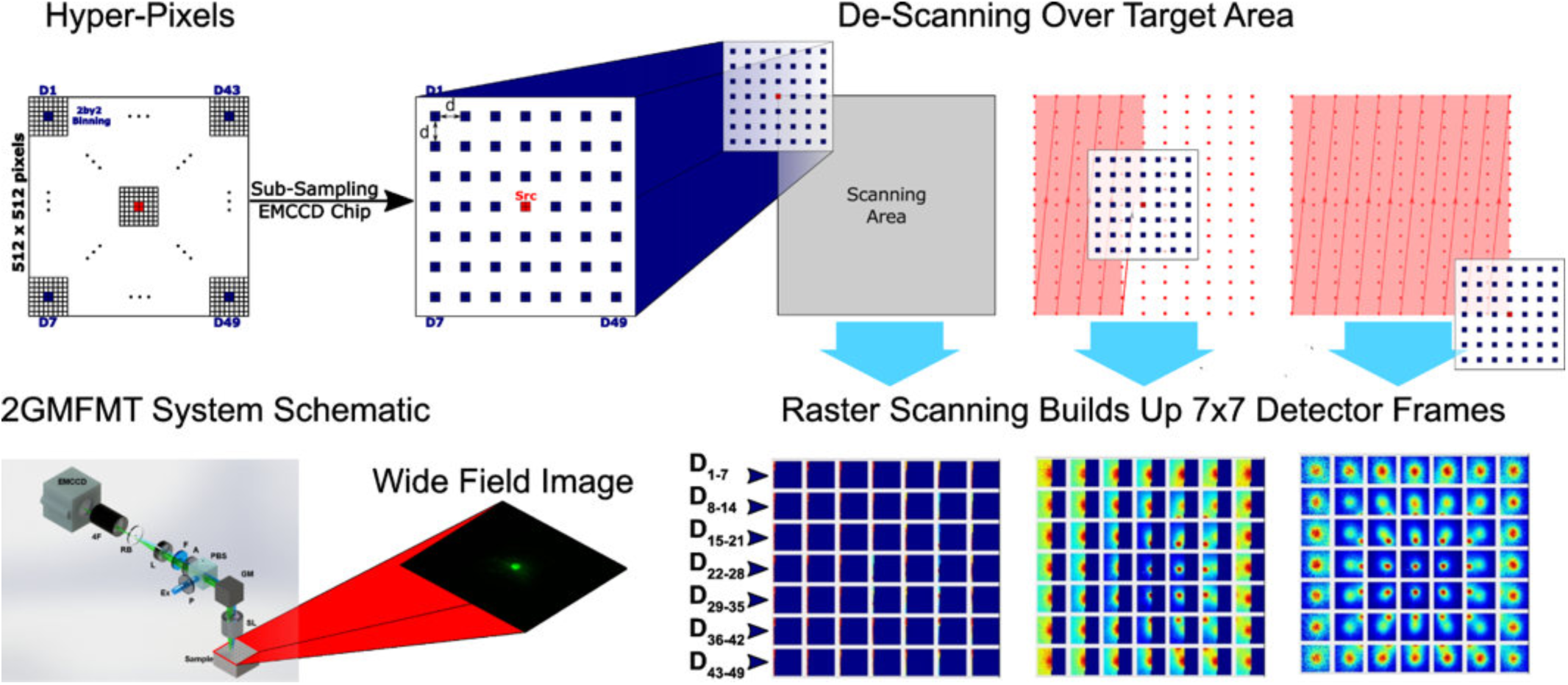
Full EMCCD chip with 2by2 binning was used for most of the acquisition schemes. Users have the flexibility adjust the source-detector separation and total detector number. De-scanning mode enables user-defined sampling density, seamlessly selecting different detector number and separations. In the descanning scheme of the 2GMFMT system, detectors collect the data in parallel. Once the raster scanning completed, full detector data set is ready.

**Supplementary Figure 4.**
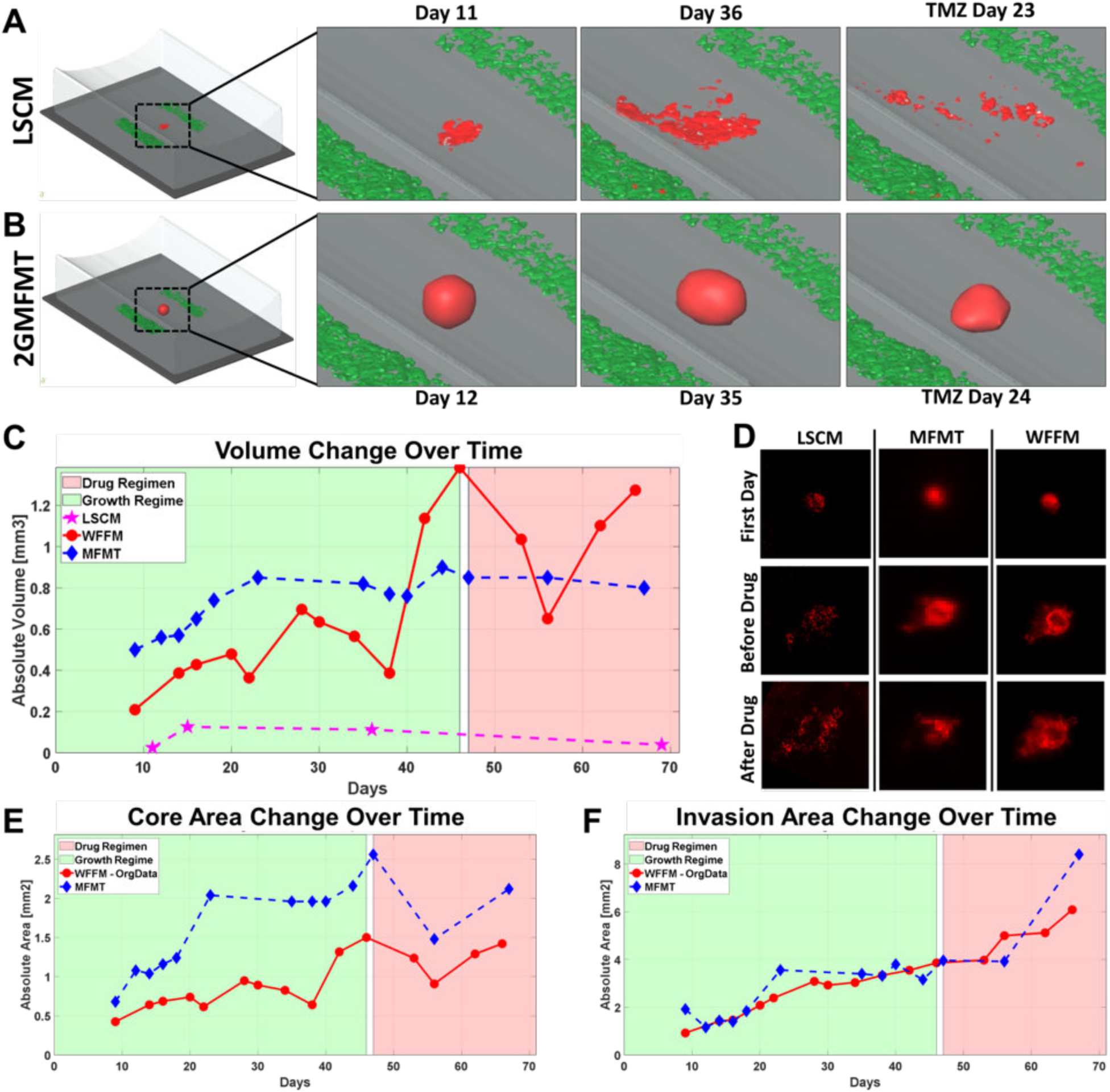
Comparison of imaging modalities monitoring tumor growth on sample 2 (Tumor ID=C). (A&B) Longitudinal volumetric assessment by LSCM (A) and 2GMFMT (B). (C) Volume change of GBM tumor mass measured by LSCM, WFFM, and MFMT. WFFM and MFMT showed similar trends overall while LSCM measurements were significantly under-estimated.(D) Maximum intensity projections from LSCM and 2GMFMT were compared with WFFM images. MFMT captured invasion patterns observed in WFFM with lower resolution while LSCM detected limited regions of the tumor invasions. (E&F) Longitudinal intensity assessment of GBM tumor comparing WFFM and MFMT. The intensity change trends in the core area (E) and the invasion area (F) showed similarities between the two modalities.

**Supplementary Figure 5.**
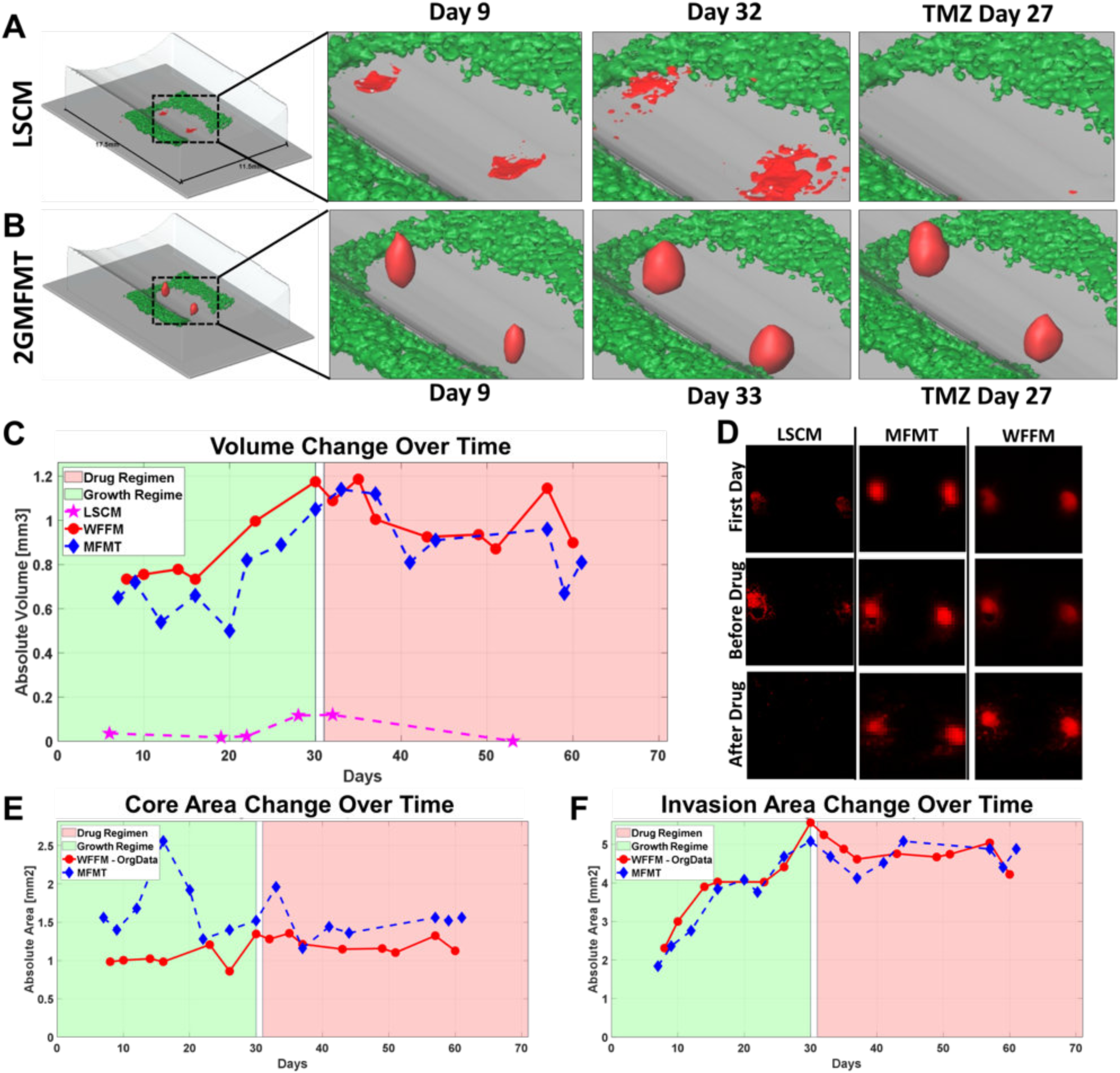
Comparison of imaging modalities monitoring tumor growth on sample 3 (Tumor ID=G). (A&B) Longitudinal volumetric assessment by LSCM (A) and 2GMFMT (B). (C) Volume change of GBM tumor mass measured by LSCM, WFFM, and MFMT. WFFM and MFMT showed similar trends overall while LSCM measurements were significantly under-estimated. (D) Maximum intensity projections from LSCM and 2GMFMT were compared with WFFM images. MFMT captured invasion patterns observed in WFFM with lower resolution while LSCM detected limited regions of the tumor invasions. (E&F) Longitudinal intensity assessment of GBM tumor comparing WFFM and MFMT. The intensity change trends in the core area (E) and the invasion area (F) showed similarities between the two modalities.

**Supplementary Figure 6.**
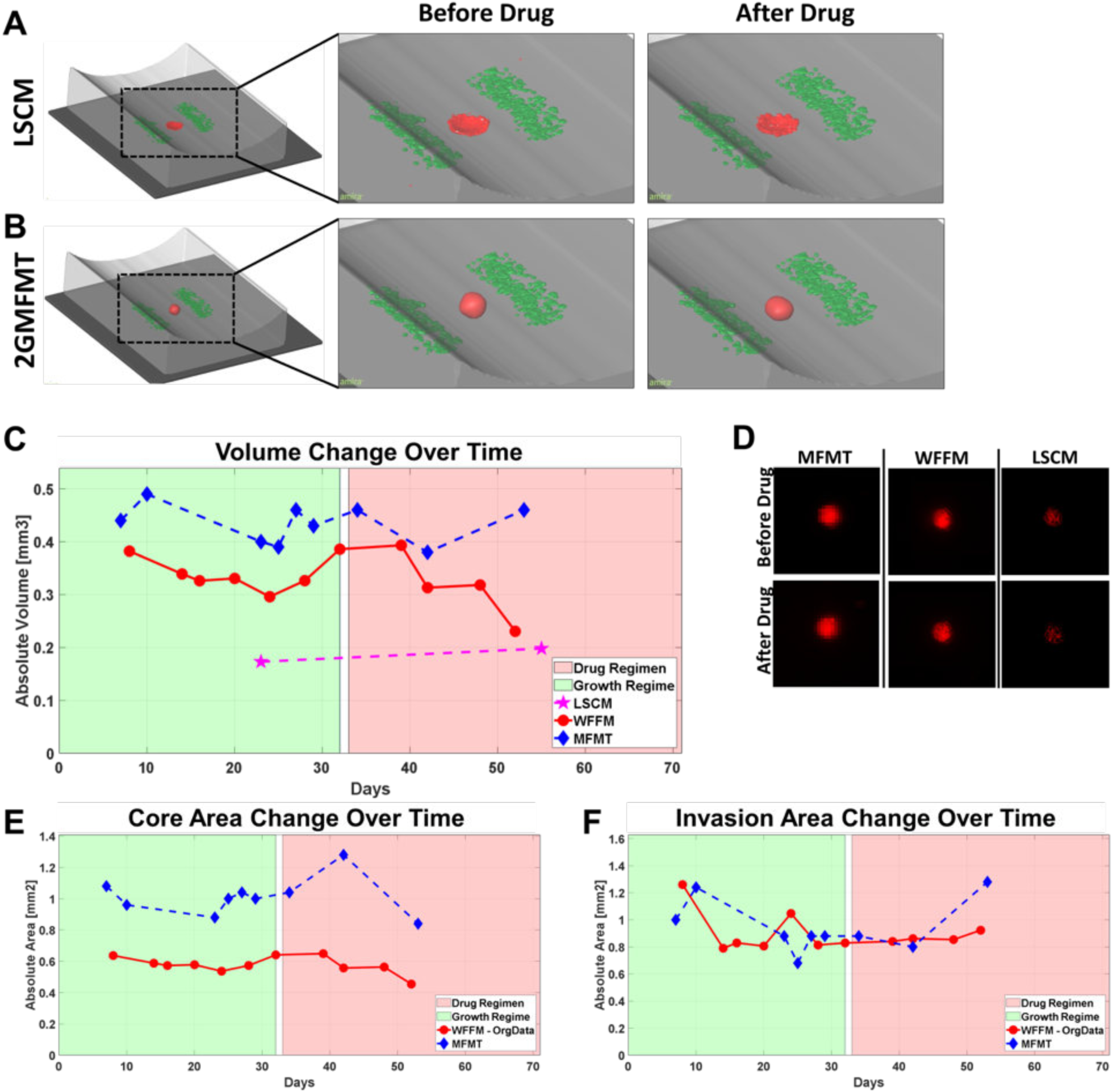
Comparison of imaging modalities monitoring tumor growth on sample 4 (Tumor ID=E). (A&B) Longitudinal volumetric assessment by LSCM (A) and 2GMFMT (B). (C) Volume change of GBM tumor mass measured by LSCM, WFFM, and MFMT. WFFM and MFMT showed similar trends overall while LSCM measurements were significantly under-estimated. (D) Maximum intensity projections from LSCM and 2GMFMT were compared with WFFM images. MFMT captured invasion patterns observed in WFFM with lower resolution while LSCM detected limited regions of the tumor invasions. (E&F) Longitudinal intensity assessment of GBM tumor comparing WFFM and MFMT. The intensity change trends in the core area (E) and the invasion area (F) showed similarities between the two modalities.

## References

[1] R. Stupp et al., “Radiotherapy plus Concomitant and Adjuvant Temozolomide for Glioblastoma,” N Engl J Med, vol. 35210, 2005.

[2] R. Stupp et al., “Effects of radiotherapy with concomitant and adjuvant temozolomide versus radiotherapy alone on survival in glioblastoma in a randomised phase III study: 5-year analysis of the EORTC-NCIC trial,” Lancet Oncol., vol. 10, no. 5, pp. 459–466, May 2009.

[3] J. D. Lathia, S. C. Mack, E. E. Mulkearns-Hubert, C. L. L. Valentim, and J. N. Rich, “Cancer stem cells in glioblastoma,” Genes Dev., vol. 29, no. 12, pp. 1203–1217, Jun. 2015.

[4] D. Hambardzumyan and G. Bergers, “Glioblastoma: Defining Tumor Niches,” Trends in Cancer, vol. 1, no. 4, pp. 252–265, Dec. 2015.

[5] R. J. Gilbertson and J. N. Rich, “Making a tumour’s bed: glioblastoma stem cells and the vascular niche.,” Nat. Rev. Cancer, vol. 7, no. 10, pp. 733–736, 2007.

[6] S. Yu et al., “Isolation and characterization of cancer stem cells from a human glioblastoma cell line U87,” Cancer Lett., vol. 265, no. 1, pp. 124–134, Jun. 2008.

[7] Y. Xie et al., “The Human Glioblastoma Cell Culture Resource: Validated Cell Models Representing All Molecular Subtypes,” EBioMedicine, vol. 2, no. 10, pp. 1351–1363, Oct. 2015.

[8] Y. Imamura et al., “Comparison of 2D- and 3D-culture models as drug-testing platforms in breast cancer,” Oncol. Rep., vol. 33, no. 4, pp. 1837–1843, Apr. 2015.

[9] K. M. Havas et al., “Metabolic shifts in residual breast cancer drive tumor recurrence,” J. Clin. Invest., vol. 127, no. 6, pp. 2091–2105, 2017.

[10] J. Friedrich, C. Seidel, R. Ebner, and L. A. Kunz-Schughart, “Spheroid-based drug screen: considerations and practical approach,” Nat. Protoc., vol. 4, no. 3, pp. 309–324, Mar. 2009.

[11] D. Fried, R. E. Glena, J. D. Featherstone, and W. Seka, “Nature of light scattering in dental enamel and dentin at visible and near-infrared wavelengths.,” Appl. Opt., vol. 34, no. 7, pp. 1278–85, Mar. 1995.

[12] S. Nath and G. R. Devi, “Three-dimensional culture systems in cancer research: Focus on tumor spheroid model,” Pharmacol. Ther., vol. 163, pp. 94–108, Jul. 2016.

[13] S. A. Langhans, “Three-Dimensional in Vitro Cell Culture Models in Drug Discovery and Drug Repositioning,” Front. Pharmacol., vol. 9, Jan. 2018.

[14] S. S. Jensen et al., “Establishment and Characterization of a Tumor Stem Cell-Based Glioblastoma Invasion Model,” PLoS One, vol. 11, no. 7, p. e0159746, Jul. 2016.

[15] J. D. Lathia et al., “Direct In Vivo Evidence for Tumor Propagation by Glioblastoma Cancer Stem Cells,” PLoS One, vol. 6, no. 9, p. e24807, Sep. 2011.

[16] B. J. Vakoc et al., “Three-dimensional microscopy of the tumor microenvironment in vivo using optical frequency domain imaging.,” Nat. Med., vol. 15, no. 10, pp. 1219–23, Oct. 2009.

[17] J. You, Q. Zhang, K. Park, C. Du, and Y. Pan, “Quantitative imaging of microvascular blood flow networks in deep cortical layers by 1310 nm μODT,” Opt. Lett., vol. 40, no. 18, p. 4293, Sep. 2015.

[18] M. S. Ozturk, V. K. Lee, L. Zhao, G. Dai, and X. Intes, “Mesoscopic fluorescence molecular tomography of reporter genes in bioprinted thick tissue.,” J. Biomed. Opt., vol. 18, no. 10, p. 100501, Oct. 2013.

[19] J. M. Kelm, N. E. Timmins, C. J. Brown, M. Fussenegger, and L. K. Nielsen, “Method for generation of homogeneous multicellular tumor spheroids applicable to a wide variety of cell types,” Biotechnol. Bioeng., vol. 83, no. 2, pp. 173–180, Jul. 2003.

[20] M. Curie, S. Bernard, and P. Cedex, “3D Processing and Analysis with ImageJ e,” 2008.

[21] S. R. Prasad, K. S. Jhaveri, S. Saini, P. F. Hahn, E. F. Halpern, and J. E. Sumner, “CT tumor measurement for therapeutic response assessment: comparison of unidimensional, bidimensional, and volumetric techniques initial observations.,” Radiology, vol. 225, no. 2, pp. 416–419, Nov. 2002.

[22] C. G. Hubert et al., “A Three-Dimensional Organoid Culture System Derived from Human Glioblastomas Recapitulates the Hypoxic Gradients and Cancer Stem Cell Heterogeneity of Tumors Found In Vivo,” Cancer Res., vol. 76, no. 8, pp. 2465–2477, Apr. 2016.

[23] D. Dutta, I. Heo, and H. Clevers, “Disease Modeling in Stem Cell-Derived 3D Organoid Systems,” Trends Mol. Med., vol. 23, no. 5, pp. 393–410, May 2017.

[24] U. Krzic, S. Gunther, T. E. Saunders, S. J. Streichan, and L. Hufnagel, “Multiview light-sheet microscope for rapid in toto imaging,” Nat. Methods, vol. 9, no. 7, pp. 730–733, Jul. 2012.

[25] V. K. Lee, A. M. Lanzi, H. Ngo, S.-S. Yoo, P. a. Vincent, and G. Dai, “Generation of Multi-scale Vascular Network System Within 3D Hydrogel Using 3D Bio-printing Technology,” Cell. Mol. Bioeng., vol. 7, no. 3, pp. 460–472, Sep. 2014.

[26] V. K. Lee et al., “Creating perfused functional vascular channels using 3D bio-printing technology.,” Biomaterials, vol. 35, no. 28, pp. 8092–102, Sep. 2014.

[27] W. Lee et al., “Multi-layered culture of human skin fibroblasts and keratinocytes through three-dimensional freeform fabrication.,” Biomaterials, vol. 30, no. 8, pp. 1587–95, Mar. 2009.

[28] W. Lee et al., “On-demand three-dimensional freeform fabrication of multi-layered hydrogel scaffold with fluidic channels.,” Biotechnol. Bioeng., vol. 105, no. 6, pp. 1178–1186, Apr. 2010.

[29] J. Chen and X. Intes, “Comparison of Monte Carlo methods for fluorescence molecular tomography— computational efficiency,” Med. Phys., vol. 38, no. 10, p. 5788, Oct. 2011.

[30] F. Yang, M. S. Ozturk, R. Yao, and X. Intes, “Improving mesoscopic fluorescence molecular tomography through data reduction,” Biomed. Opt. Express, vol. 8, no. 8, pp. 3868–3881, Aug. 2017.

[31] R. Favicchio et al., “Quantitative performance characterization of three-dimensional noncontact fluorescence molecular tomography,” J. Biomed. Opt., vol. 21, no. 2, p. 026009, Feb. 2016.

[32] L. Zhao, H. Yang, W. Cong, G. Wang, and X. Intes, “L_p regularization for early gate fluorescence molecular tomography,” Opt. Lett., vol. 39, no. 14, p. 4156, Jul. 2014.

[33] X. Intes, J. Ripoll, Y. Chen, S. Nioka, A. G. Yodh, and B. Chance, “In vivo continuous-wave optical breast imaging enhanced with Indocyanine Green,” Med. Phys., vol. 30, no. 6, p. 1039, May 2003.

